# Microbial Metabolites Potentiate MAIT Cell Anti-Tumor Immunity Against Solid Tumors

**DOI:** 10.64898/2026.07.24.740456

**Authors:** Yichen Zhu, Xinyuan Shen, Yuning Chen, Nathan Ma, Catherine Zhang, Annabel S. Zhao, Yanxin Tian, Shreya Gumate, Jie Huang, Siyu Lin, Aijun Wang, Vatche G. Agopian, Yan-Ruide Li, Lili Yang

## Abstract

Mucosal-associated invariant T (MAIT) cells sense riboflavin metabolites through the monomorphic antigen-presenting molecule MR1, providing a unique opportunity to therapeutically mobilize a broadly shared T cell population without genetic engineering. Here, we show that the highly potent microbial metabolites, 5-OP-RU and 5-OE-RU, can be exploited as pharmacologic precision immune activators to drive human MAIT cell responses against solid tumors. Ligand stimulation in human co-culture systems elicited robust MAIT cell cytotoxicity, inflammatory cytokine secretion, and transcriptional states transformation. *In vivo* administration of riboflavin ligands 5-OP-RU significantly suppressed tumor growth in xenograft liver cancer models. Metabolite-driven MAIT activation also reprogrammed the local immune landscape, enhancing effector function and overcoming features of the immunosuppressive niche by markedly eliminating tumor-associated macrophages (TAMs) and myeloid-derived suppressor cells (MDSCs) within the tumor microenvironment (TME). These findings reveal that microbial riboflavin metabolites can power and redirect MAIT cells to solid tumors, establishing MR1–metabolite signaling as a tractable therapeutic axis for liver cancer and other solid malignancies.

## Introduction

Mucosal-associated invariant T (MAIT) cells are a conserved subset of unconventional αβ T cells with rapid effector capacity and broad antimicrobial and immunoregulatory activity^1–8^. MAIT cells express a semi-invariant T cell receptor that recognizes the monomorphic antigen-presenting molecule MR1 loaded with small-molecule ligands derived from microbial riboflavin biosynthesis, establishing an antigen-recognition axis that is distinct from peptide–HLA–restricted immunity^1–5^. Although MAIT cells typically comprise only a minor fraction of circulating T cells, they are highly enriched in mucosal and barrier-associated tissues, where they can represent a substantial proportion of resident lymphocytes, particularly in organs such as the liver, lung, and intestine^3,4,9^. This tissue distribution, together with their innate-like transcriptional program, positions MAIT cells as sentinels tuned to metabolite sensing in peripheral tissues.

MR1-restricted immunity has recently gained prominence as an attractive framework for population-wide cancer targeting^10,11^. MR1 is monomorphic and broadly expressed, and MR1-dependent recognition of transformed cells has been implicated as a shared vulnerability across diverse malignancies, motivating interest in MR1 as a pan-population antigen-presenting platform^10^. MAIT cells are detected within human tumors and adjacent tissues across cancer types, consistent with their ability to access barrier and visceral sites and their intrinsic cytotoxic potential^12–20^. However, the net contribution of MAIT cells to tumor immunity remains context dependent and frequently constrained by the TME^16–19,21^. Tumor-infiltrating MAIT cells in multiple solid tumors have been reported to exhibit diminished cytotoxicity and altered cytokine profiles accompanied by increased expression of inhibitory receptors and activation markers, indicating chronic stimulation coupled to functional impairment^12–15^. Suppressive myeloid circuits, including tumor-associated macrophages and related immunosuppressive populations, recur as key correlates of MAIT dysfunction, suggesting that MAIT cells may be present but inadequately licensed to execute effective antitumor programs within solid tumors^13^. These observations collectively support a model in which MAIT cells represent a potentially powerful effector compartment whose antitumor activity is limited by insufficient productive MR1–ligand signaling and by dominant myeloid-mediated immune suppression.

Microbial riboflavin metabolites constitute the dominant class of physiological MR1 ligands capable of triggering MAIT TCR signaling and programming MAIT effector function^4,5,22,23^. This axis is central to MAIT cell development and maturation, and microbial community composition can influence the magnitude and quality of MAIT activation through differential metabolite production^24–26^. Riboflavin-derived ligands such as 5-OP-RU therefore provide a mechanistically direct route to engage MAIT cells through their defining antigen receptor, enabling antigen-specific activation without genetic engineering^4,22^. In infection and inflammatory contexts, MR1 ligands can drive MAIT activation, cytokine production, and cytotoxic responses, supporting the concept that small-molecule agonism can tune MAIT functional state^27–30^. In cancer, preclinical studies in mice have shown that MAIT activation can be induced *in vivo* under conditions of strong innate stimulation, and that this activation can associate with antitumor effects in multiple tumor models^31,32^. However, mechanistic interpretation and translational extrapolation are complicated by fundamental interspecies differences: murine MAIT cells are sparse compared with humans, are skewed toward distinct functional subsets, and often require adjuvant-driven inflammatory cues for robust expansion and activation^3,29,33,34^. In contrast, human MAIT cells are abundant in relevant tissues and exhibit broader functional plasticity, with mixed type-1 and type-17 programs and measurable cytotoxic potential, emphasizing the need for direct investigation of microbial metabolite–driven MAIT responses in human systems^4,33,34^.

Here, we establish a framework for MAIT cell activation and targeting of solid tumors by leveraging microbial riboflavin metabolites as precision immune agonists. We demonstrate that microbial metabolites selectively mobilize and expand MAIT cells across both healthy donors and cancer patients, enabling broad MR1-dependent MAIT cytotoxicity against diverse solid tumor types, including liver, ovarian, melanoma, lung, breast, and colorectal cancers. Microbial metabolite–driven activation induces coordinated effector programs in MAIT cells characterized by enhanced cytotoxic capacity and inflammatory cytokine production, consistent with the rapid effector responses typical of innate-like T cell populations. Beyond direct tumor cell killing, we further identify a dual targeting capability whereby microbial metabolite–activated MAIT cells preferentially eliminate MR1-expressing immunosuppressive myeloid populations within the TME. Together, these findings position microbial metabolites as tractable pharmacologic activators of human MAIT cells and establish MR1–microbial metabolite signaling as a therapeutic axis for mobilizing MAIT-mediated immunity against solid tumors.

## Results

### MAIT cells exhibit a distinct effector-memory phenotype and intrinsic cytotoxic program compared to conventional T cells

To establish a foundational understanding of the intrinsic properties of human MAIT cells relevant to therapeutic exploitation, we first investigated their distribution and phenotypic characteristics relative to conventional T cell subsets. Based on previous studies, human MAIT cells exhibit distinct tissue biodistributions; although they comprise ∼2% of the total T cell population in peripheral blood, they represent a predominant T cell subtype in mucosal tissues including the lung, liver, and intestine (Fig. 1a&1b)^21,35–38^. MAIT cells were identified with MR1/5-OP-RU tetramer and confirmed by high expression of TCR Vα7.2 and CD161 (Fig. 1c). Phenotypic comparison of MAIT cells with conventional CD4⁺ and CD8⁺ T cells isolated from healthy donor blood revealed that MAIT cells are predominantly CD8⁺ or CD4⁻CD8⁻ double-negative (Fig. 1c) and display a consistent effector memory (Tem) phenotype (Fig. 1d). In contrast, circulating conventional CD4⁺ and CD8⁺ T cells comprised heterogeneous naïve (CD62L⁺CD45RO⁻), central memory (CD62L⁺CD45RO⁺), effector (CD62L⁻CD45RO⁻), and effector memory subsets, highlighting the relative functional uniformity of MAIT cells.

**Fig. 1.**
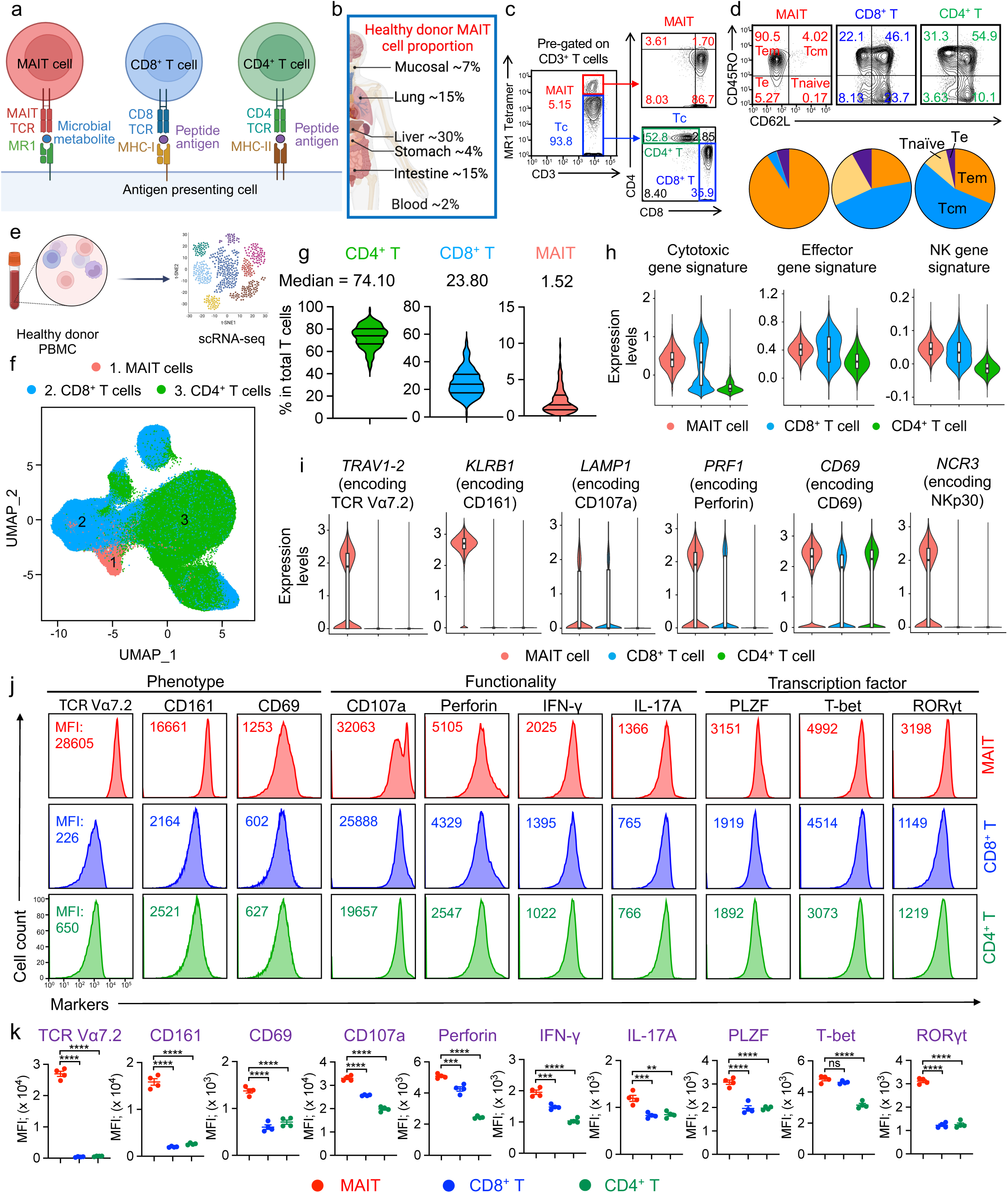
MAIT cells exhibit a distinct effector-memory phenotype and intrinsic cytotoxic program compared to conventional T cells. a. Schematics showing antigen recognition by MAIT, CD8⁺ T, and CD4⁺ T cells. b. Tissue distribution of MAIT cells in healthy humans, summarized from the published database. c. FACS detection of MAIT (CD3^+^MR1-Tetramer⁺) and Tc (CD3^+^MR1-Tetramer^-^) cells, as well as their CD4⁺ vs CD8⁺ subsets. d. FACS analysis of effector/memory phenotype based on CD45RO and CD62L expression. Tem: effector memory T cells; Tcm, central memory T cells; Tnaive: naïve T cells; Te: effector T cells. e-h. Comparison of human MAIT and conventional CD4+ and CD8+ T cells isolated from healthy donor peripheral blood using scRNA-seq analysis. e. Schematic showing the gene profiling of T cells in in healthy donor PBMCs. f. Combined UMAP plot showing the identification of MAIT cells. Each dot represents a single cell and is colored according to its cell cluster assignment. g. Proportion of MAIT cell subset among total T cells. h-i. scRNA-seq analysis of the expression of the indicated gene signatures (h) and genes (i) in MAIT cells. j. FACS detection of surface markers, intracellular cytokines, cytotoxic molecules and transcription factors in MAIT cells. CD4^+^ T and CD8^+^ T cells were included as controls. k. Quantification of j. Representative of 1 (e-i) and 3 (c-d, j-k) experiments. Data are presented as the mean ± SEM. **p < 0.01, ***p < 0.001, ****p < 0.0001 by one-way ANOVA (k).

To further define the molecular features underlying this distinct phenotype, we analyzed a publicly available single-cell RNA sequencing dataset of human PBMC samples derived from 166 donors (Fig. 1e)^39^. Within this dataset, MAIT cells were identified by co-expression of *TRAV1-2* (encoding TCR Vα7.2) and *KLRB1* (encoding CD161) (Fig. 1f&1i). MAIT cells were consistently detected across donors, with a median frequency of 1.52% among total T cells, although inter-individual variability was observed, with some donors exceeding 10% (Fig. 1g). Transcriptomic analysis revealed that MAIT cells exhibited elevated expression of cytotoxic, effector, and natural killer (NK)-associated programs relative to conventional T cells, including genes encoding CD107a (*LAMP1*), perforin (*PRF1*), CD69, and NKp30 (*NCR3*) (Fig. 1h&1i). These findings suggest that circulating MAIT cells possess a pre-armed effector state characterized by cytotoxic and innate-like functional features.

We next validated these observations at the protein level using flow cytometry of healthy donor PBMC samples (Fig. 1j&1k). Consistent with transcriptomic findings, MAIT cells demonstrated increased expression of cytotoxic mediators (CD107a and perforin) and activation marker CD69. In addition, MAIT cells produced IFN-γ and IL-17A and expressed lineage-defining transcription factors including PLZF, which regulates innate-like T cell development, T-bet, the driver of Th1 effector responses, and RORγt, the canonical regulator of Th17 programs (Fig. 1j–k), supporting a multifunctional effector profile.

Collectively, these data establish that human circulating MAIT cells represent a functionally poised T cell population characterized by effector memory bias, intrinsic cytotoxic capacity, and innate-like transcriptional programming. This combination of features provides a strong biological rationale for exploring strategies to therapeutically mobilize MAIT cells for cancer immunotherapy.

### Microbial metabolites selectively mobilize and expand human MAIT cells across healthy donors and cancer patients

Given that circulating MAIT cells exhibit intrinsic cytotoxic and effector properties yet remain relatively infrequent in peripheral blood, we next sought to determine whether pharmacologic activation through the MAIT TCR could selectively mobilize this population. Because microbial riboflavin metabolites are naturally presented by MR1 and trigger MAIT TCR signaling, we hypothesized that such ligands could function as precision immune activators capable of inducing MAIT cell expansion and activation within physiologic immune cell contexts^40–43^.

To test this hypothesis, we evaluated the capacity of the potent MR1 ligand 5-OP-RU to activate MAIT cells in PBMCs derived from healthy donors and cancer patients (Fig. 2a). Notably, PBMC cultures contain endogenous antigen-presenting cells (APCs) capable of MR1-mediated ligand presentation, thereby enabling physiologic activation conditions. Following addition of 5-OP-RU to PBMC cultures, robust MAIT cell enrichment was observed after 7 days (Fig. 2b). Although MAIT cells comprised less than 10% of total CD3⁺ T cells at baseline (Fig. 1g), microbial metabolite stimulation increased their frequency to as high as ∼60% in healthy donor samples (Fig. 2b), indicating strong selective expansion.

**Fig. 2.**
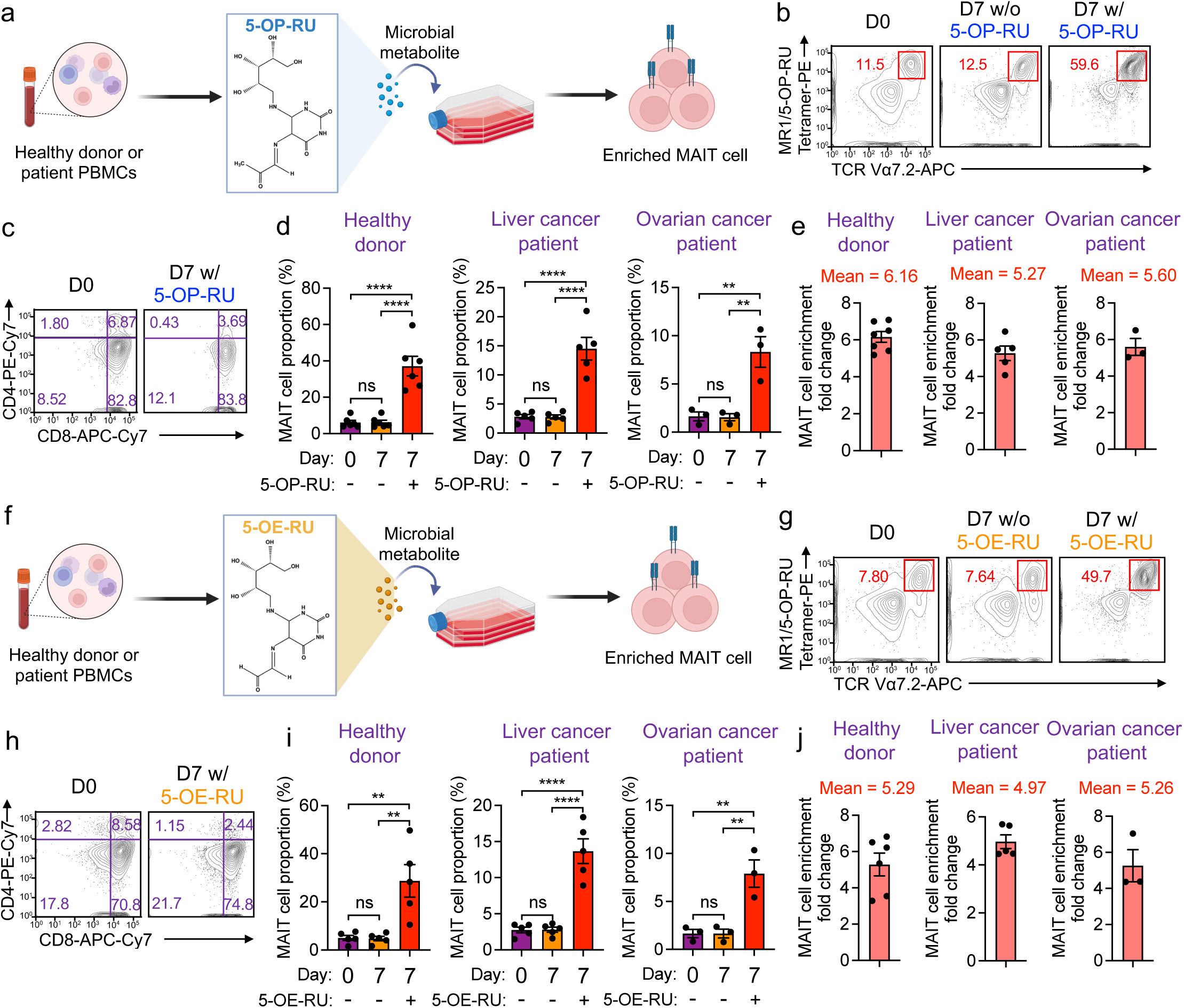
Microbial metabolites selectively mobilize and expand human MAIT cells across healthy donors and cancer patients. a. Schematic for riboflavin-derived microbial metabolite 5-OP-RU stimulation of MAIT cells in PBMCs from healthy donors or cancer patients. b. FACS plots showing MR1/5-OP-RU tetramer⁺ TCR Vα7.2⁺ MAIT cells at baseline (D0), day 7 without 5-OP-RU, and day 7 with 5-OP-RU stimulation. c. CD4 and CD8 subset distribution of MAIT cells at D0 and D7 with 5-OP-RU stimulation. d. Quantification of MAIT cell proportions in PBMCs from healthy donors (n = 6), liver cancer patients (n = 5), and ovarian cancer patients (n = 3) at D0 and D7 ± 5-OP-RU stimulation. e. Fold enrichment of MAIT cells after 5-OP-RU stimulation in healthy donors (n = 6), liver cancer patients (n = 5), and ovarian cancer patients (n = 3). f. Schematic for riboflavin-derived microbial metabolite 5-OE-RU stimulation of MAIT cells in PBMCs from healthy donors or cancer patients. g. FACS plots showing MR1/5-OP-RU tetramer⁺ TCR Vα7.2⁺ MAIT cells at baseline (D0), day 7 without 5-OE-RU, and day 7 with 5-OE-RU stimulation. h. CD4 and CD8 subset distribution of MAIT cells at D0 and D7 with 5-OE-RU stimulation. i. Quantification of MAIT cell proportions in PBMCs from healthy donors (n = 6), liver cancer patients (n = 5), and ovarian cancer patients (n = 3) at D0 and D7 ± 5-OE-RU stimulation. j. Fold enrichment of MAIT cells after 5-OE-RU stimulation in healthy donors (n = 6), liver cancer patients (n = 5), and ovarian cancer patients (n = 3). Representative of 3 experiments. Data are presented as the mean ± SEM. ns, not significant, **p < 0.01, ****p < 0.0001 by one-way ANOVA (d, i). See also Supplementary Fig. 1.

We next examined whether this phenomenon extended to cancer patient samples. PBMCs from five liver cancer patients and three ovarian cancer patients displayed heterogeneous baseline MAIT cell frequencies; however, 5-OP-RU consistently induced marked MAIT cell enrichment across all donors (Fig. 2c&2d). Greater than five-fold increases in MAIT cell frequency was observed following microbial metabolite stimulation in both healthy and cancer patient cohorts, demonstrating the robustness of this activation strategy across distinct clinical backgrounds (Fig. 2e).

To determine whether this effect was specific to 5-OP-RU, we performed parallel experiments using an additional riboflavin-derived MR1 ligand, 5-OE-RU, which exhibits lower potency in activating MAIT TCR signaling (Fig. 2f)^44–46^. Similar stimulation assays revealed that 5-OE-RU also promoted MAIT cell enrichment and expansion after 7 days (Fig. 2g&2h). Although the magnitude of expansion was modestly reduced relative to 5-OP-RU, significant increases in MAIT cell frequency were observed across healthy donors and cancer patients (Fig. 2i&2j), indicating that MAIT cell mobilization can be achieved across a range of MR1 ligands.

Collectively, these findings demonstrate that microbial riboflavin metabolites can selectively enrich and expand MAIT cells within human PBMC environments from both healthy individuals and cancer patients. These results provide functional evidence that metabolite-mediated MAIT activation represents a pharmacologic strategy capable of mobilizing endogenous MAIT cells, supporting further investigation of MR1–microbial metabolite signaling as a tractable immune activation axis for solid tumor immunotherapy.

### Microbial metabolite-driven MR1 signaling enables broad MAIT cell cytotoxicity across solid tumors

To enable functional interrogation of MAIT cell anti-tumor activity, MAIT cells were generated from healthy donors and cancer patients using MR1/5-OP-RU tetramer–based sorting followed by expansion with irradiated PBMC feeder cells pulsed with 5-OP-RU, yielding highly purified MAIT cell populations suitable for downstream assays (Supplementary Fig. 1).

We next sought to determine whether microbial metabolite–mediated MAIT activation could translate into tumor-directed cytotoxic function across diverse solid malignancies. To this end, *in vitro* killing assays were performed against multiple cancer types, including liver, ovarian, melanoma, lung, breast, and colorectal cancers (Fig. 3a). Twelve human cancer cell lines representing distinct genetic backgrounds and tissue origins were evaluated (Supplementary Fig. 2a). Ten parental tumor cell lines, including liver cancer cell line HepG2, Hep3B, SNU423, and SK-HEP-1, ovarian cancer cell line OVCAR3m and SKOV3, melanoma cell line A375, lung cancer cell ling H226, breast cancer cell line MDA-MB-231, and colorectal cancer cell line HCT116) were engineered to express a firefly luciferase–EGFP dual reporter (FG), enabling quantitative assessment of tumor cell viability by luminescence and flow cytometry (Supplementary Fig. 2a). To directly interrogate MR1 dependency, SK-HEP-1 cells were further engineered to overexpress MR1 (SK-HEP-1-MR1-FG) or lack MR1 (SK-HEP-1-FG^MR1−/−^) (Supplementary Fig. 2a). Baseline characterization revealed heterogeneous MR1 expression across the tumor panel, indicating that MR1 is broadly present across human solid tumors and may serve as a shared molecular platform for MAIT cell–mediated tumor recognition^10,47–50^ (Fig. 3b).

**Fig. 3.**
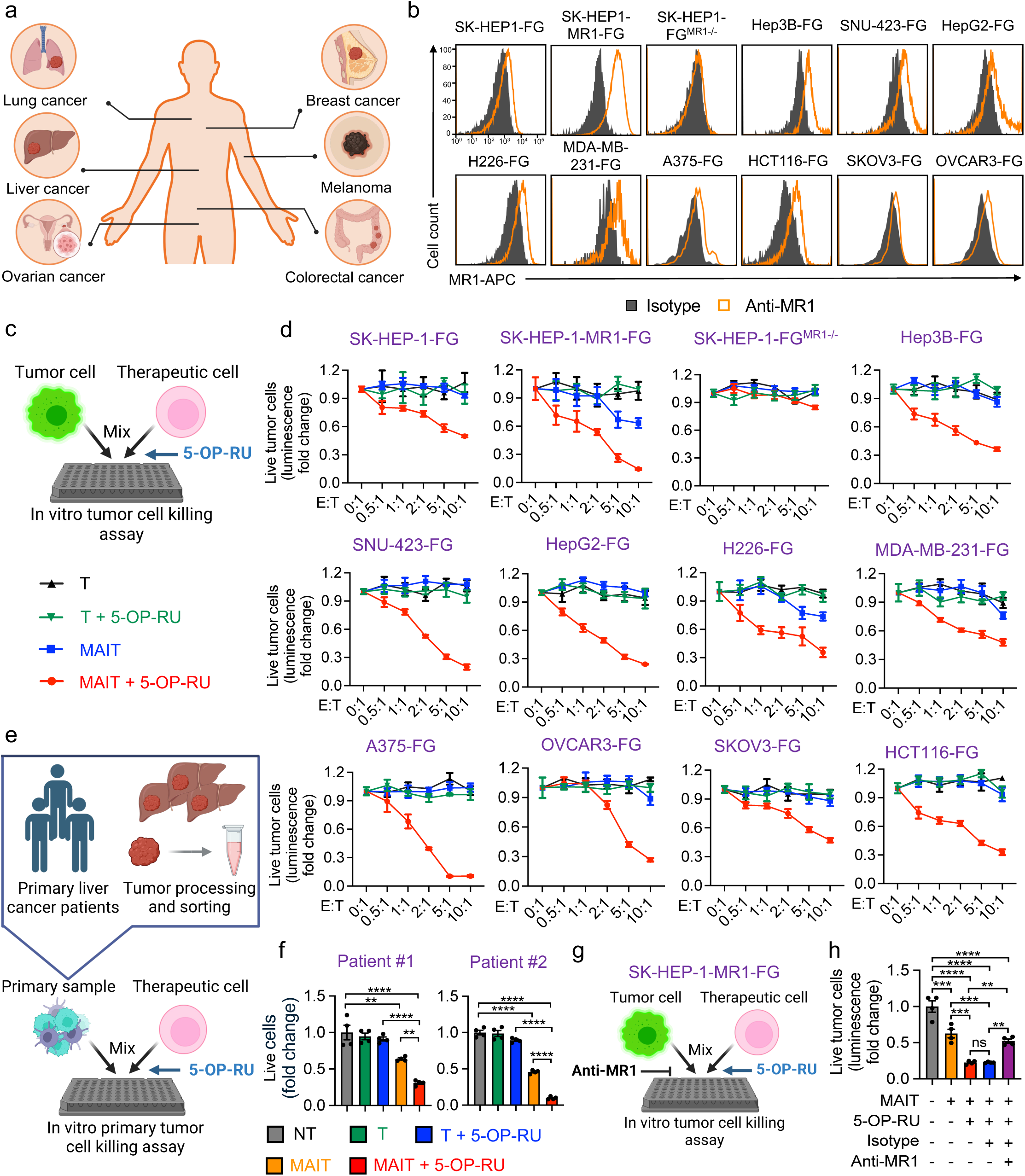
Microbial metabolite-driven MR1 signaling enables broad MAIT cell cytotoxicity across solid tumors. a. Schematics illustrating solid cancer types utilized in this study. b-d. Studying microbial metabolite-enhanced MAIT cell antitumor activity using human solid tumor cell lines. Tumor cells were cocultured with MAIT cells in the presence or absence of 5-OP-RU. T cells were included as a therapeutic cell control. b. FACS detection of MR1 expression on the indicated tumor cells. c. Experimental design. d. Tumor cell killing data at 24 h (n = 4). e-f. Studying microbial metabolites-enhanced MAIT cell antitumor activity using primary liver cancer patient samples. Tumor cells were cocultured with MAIT cells in the presence or absence of 5-OP-RU. e. Experimental design. f. Primary liver tumor cell killing data at 24 h (n = 4). g-h. Studying the tumor cell killing mechanisms of MAIT cells mediated by MR1 and microbial metabolite, 5-OP-RU. g. Experimental design. h. Tumor cell killing data at 24 h (E:T ratio = 5:1; n = 4). Representative of 3 experiments. Data are presented as the mean ± SEM. ns, not significant, **p < 0.01, ***p < 0.001, ****p < 0.0001 by one-way ANOVA (f, h). See also Supplementary Fig. 2 and 3.

Healthy donor–derived MAIT or conventional T cells were co-cultured with tumor cells in the presence or absence of the microbial metabolite 5-OP-RU. Across all tumor models tested, 5-OP-RU stimulation markedly enhanced MAIT cell–mediated cytotoxicity compared with unstimulated MAIT cells (Fig. 3c&3d). In contrast, conventional T cells displayed minimal cytotoxic activity regardless of metabolite exposure, indicating that the observed anti-tumor effects were specific to MR1-restricted MAIT cell activation (Fig. 3d). Importantly, MR1-overexpressing SK-HEP-1-MR1-FG cells exhibited increased susceptibility to MAIT-mediated killing relative to parental SK-HEP-1-FG cells, whereas MR1-deficient SK-HEP-1-FG^MR1−/−^ cells demonstrated minimal killing independent of 5-OP-RU stimulation (Fig. 3d). These findings establish a direct relationship between MR1 expression and MAIT cell cytotoxic activity.

Additionally, MAIT cells alone exhibited negligible tumor killing across the cancer cell panel, indicating limited basal cytotoxic activity in the absence of exogenous ligand (Fig. 3d). The only exception was MR1-overexpressing SK-HEP-1-MR1-FG cells, which displayed modest ligand-independent killing at high effector-to-target ratios, consistent with elevated MR1 surface expression (Fig. 3d). This observation is consistent with reports that MAIT cells are detectable within many solid TME, suggesting that insufficient MR1–ligand availability represents a key constraint on their effector function *in situ*^13,15,51^.

To determine whether this response generalized across MR1 ligands, parallel experiments were conducted using the riboflavin-derived metabolite 5-OE-RU. Consistent with 5-OP-RU findings, 5-OE-RU stimulation enhanced MAIT cell–mediated cytotoxicity across liver cancer models (Supplementary Fig. 2b&2c). To further evaluate physiologic ligand sources, MAIT activation assays were performed using *E. coli* culture supernatants, a natural producer of riboflavin-derived MR1 ligands^25,26,52^. Bacterial supernatants similarly enhanced MAIT cell cytotoxicity, supporting the concept that endogenous microbial metabolites can potentiate MAIT anti-tumor function (Supplementary Fig. 2d&2e).

We next assessed whether these observations extended to primary human tumors. MAIT cells were co-cultured with primary liver tumor cells derived from patients (Fig. 3e). Primary tumor cells displayed variable but generally detectable MR1 expression (Supplementary Fig. 3a). Upon 5-OP-RU stimulation, MAIT cells consistently mediated enhanced cytotoxicity against patient-derived tumor cells, whereas conventional T cells exhibited minimal killing under both conditions (Fig. 3f and Supplementary Fig. 3b). These results support selective activation of MAIT cells as the dominant mechanism driving metabolite-enhanced tumor cell elimination.

Finally, MR1 blocking experiments confirmed mechanistic MR1 dependency, as antibody-mediated MR1 blockade abrogated 5-OP-RU–enhanced tumor killing (Fig. 3g&3h). Comparable results were observed with 5-OE-RU stimulation (Supplementary Fig. 2f&2g), demonstrating that MR1–ligand interactions represent the central mechanism underlying metabolite-driven MAIT tumor cytotoxicity.

Collectively, these data demonstrate that microbial riboflavin metabolites broadly potentiate MR1-dependent MAIT cell cytotoxicity across diverse solid tumor contexts, providing functional evidence that metabolite-mediated activation constitutes a pharmacologic strategy capable of activating endogenous MAIT cells for tumor targeting.

### Microbial metabolites induce MAIT cytotoxic and inflammatory effector responses

Having established that microbial metabolite stimulation enhances MAIT cell–mediated tumor killing, we next sought to define the cellular activation programs underlying this functional response. MAIT cells were co-cultured with tumor targets in the presence or absence of the microbial metabolite 5-OP-RU, followed by phenotypic and secretome profiling (Fig. 4a&4b).

**Fig. 4.**
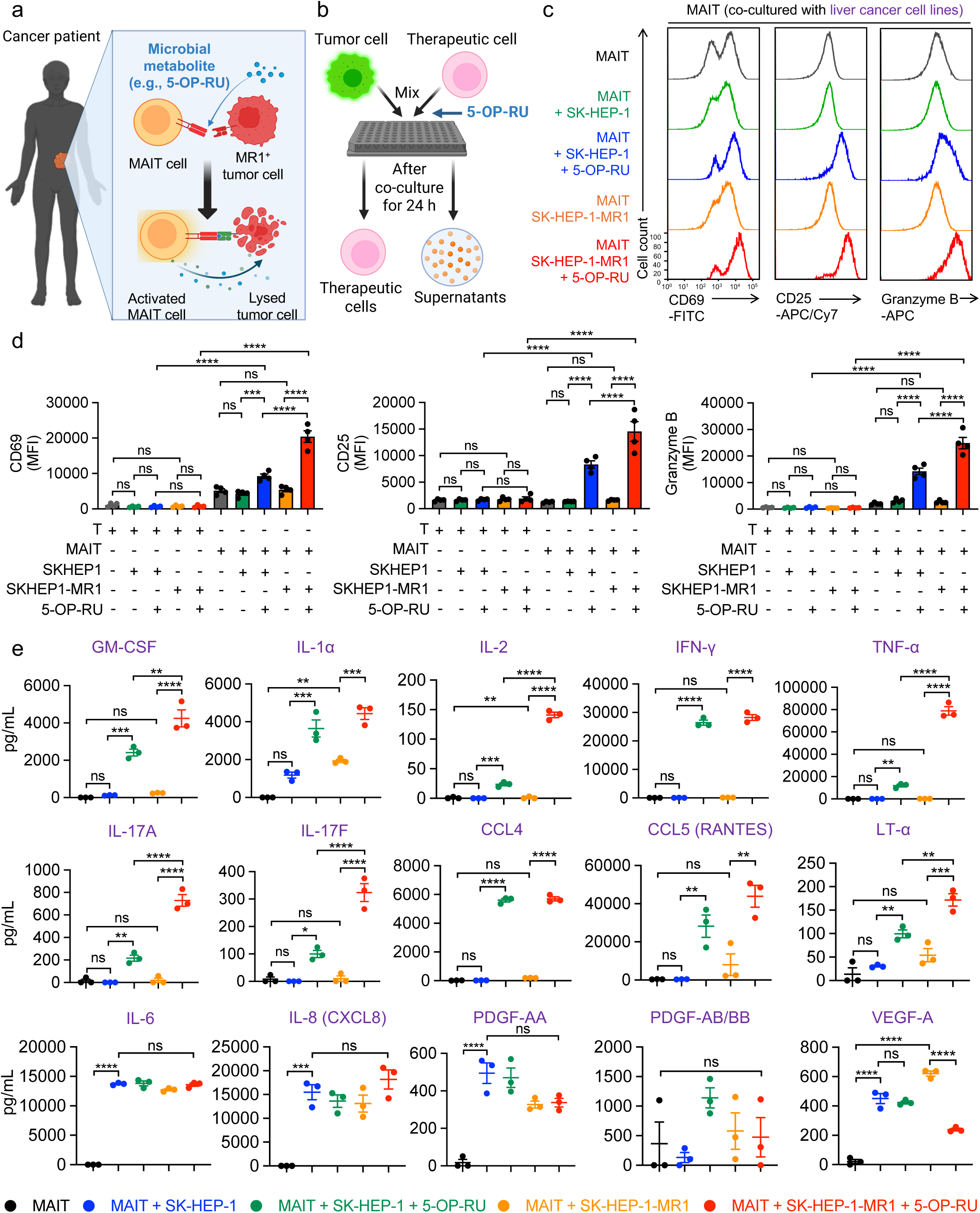
Microbial metabolites induce MAIT cytotoxic and inflammatory effector responses. a. Schematics illustrating the mechanism of MAIT cell–mediated tumor killing in cancer patients. b-e. Studying the expression of effector molecules in MAIT cells cocultured with liver cancer cells and microbial metabolite. Tumor cells were cocultured with MAIT cells in the presence or absence of 5-OP-RU. b. Experimental design. c. FACS detection of surface CD69 and CD25 expression as well as intracellular Granzyme B production of MAIT cells under the indicated conditions. d. Quantification of c (n = 4). e. Luminex analyses of cytokine and chemokine production by MAIT cells under the indicated conditions (n = 3). Representative of 1 (e) and 3 (b-d) experiments. Data are presented as the mean ± SEM. ns, not significant, **p < 0.01, ***p < 0.001, ****p < 0.0001 by one-way ANOVA (d and e). See also Supplementary Fig. 4.

Flow cytometric analysis revealed that co-culture of MAIT cells with tumor cells alone induced minimal activation, consistent with limited endogenous ligand availability (Fig. 4c&4d). In contrast, 5-OP-RU stimulation markedly increased expression of canonical activation and effector markers, including CD69, CD25, and Granzyme B (Fig. 4c&4d). This activation pattern paralleled functional cytotoxicity outcomes observed in killing assays, indicating that microbial metabolite-driven signaling directly programs MAIT effector function (Fig. 4c&4d). Notably, MR1-overexpressing SK-HEP-1-MR1-FG cells elicited stronger MAIT activation compared with parental SK-HEP-1 cells, further supporting MR1-dependent activation thresholds (Fig. 4c&4d).

To determine whether MAIT activation was accompanied by coordinated immune signaling, coculture supernatants were subjected to multiplex cytokine profiling (Fig. 4b). Metabolite-stimulated MAIT cells exhibited broad induction of inflammatory and immune-stimulatory mediators, including GM-CSF, IL-1α, IL-2, IFN-γ, TNF-α, IL-17A, IL-17F, CCL4, CCL5 (RANTES), and LT-α (Fig. 4e). These cytokines collectively reflect diverse functional axes of MAIT immunity, encompassing effector T cell activation (IL-2), cytotoxic and Th1 responses (IFN-γ, TNF-α), innate-like inflammatory signaling (GM-CSF, IL-1α), type-17 immunity (IL-17A/F), and chemokine-mediated immune cell recruitment (CCL4/CCL5)^53–57^. In contrast, mediators commonly associated with tumor-promoting or angiogenic programs, including IL-6, IL-8 (CXCL8), PDGF-AA, PDGF-AB/BB, and VEGF-A, were not induced following metabolite stimulation, suggesting selective activation of anti-tumor inflammatory pathways rather than generalized cytokine release (Fig. 4e)^58–61^.

Parallel experiments using 5-OE-RU and *E. coli* culture supernatants produced comparable activation patterns, indicating that diverse riboflavin-derived metabolites can engage a conserved MAIT activation program (Supplementary Fig. 4).

Collectively, these findings demonstrate that MR1–metabolite signaling drives coordinated MAIT cytotoxic activation and inflammatory cytokine secretion while avoiding induction of tumor-supportive mediators. These data provide mechanistic evidence that microbial metabolites function as pharmacologic immune agonists capable of reprogramming MAIT cells toward anti-tumor effector states.

### Pharmacologic MR1 activation enables MAIT cell–mediated tumor control *in vivo*

Having established that microbial metabolites drive MAIT cytotoxic activation *in vitro*, we next evaluated whether systemic delivery of microbial metabolite could potentiate MAIT cell–mediated tumor control *in vivo*. To model disseminated liver tumor growth, an orthotopic xenograft system was established by intravenous injection of SK-HEP-1-MR1-FG cells into NSG mice, enabling preferential hepatic tumor engraftment and longitudinal monitoring by bioluminescence imaging (BLI) (Fig. 5a).

**Fig. 5.**
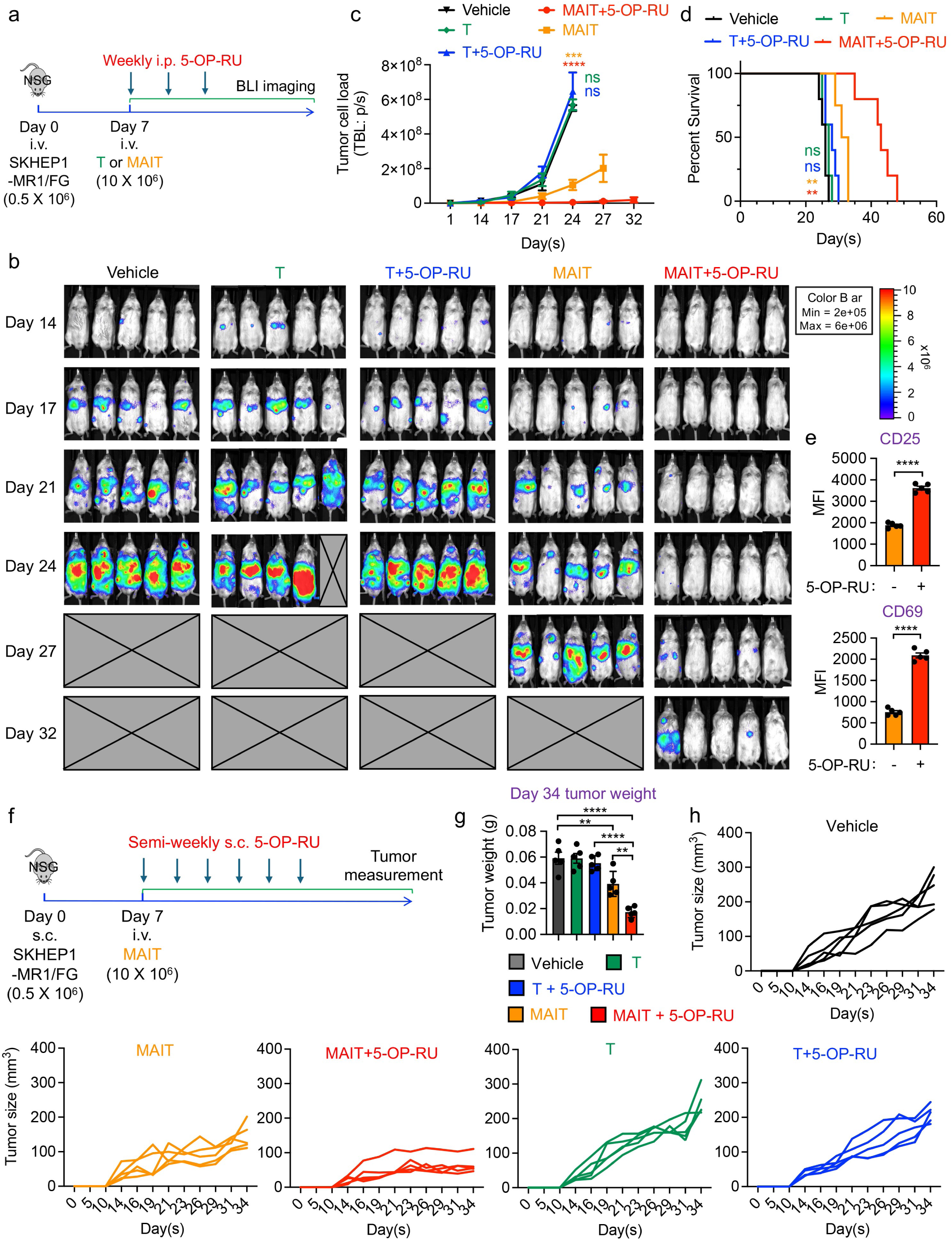
Pharmacologic MR1 activation enables MAIT cell–mediated tumor control *in vivo*. a-e. Studying the *in vivo* antitumor efficacy of MAIT cells in an orthotopic liver cancer xenograft NSG mouse model. 5-OP-RU was administrated to experimental mice weekly. a. Experimental design. BLI, live animal bioluminescence imaging; i.p., intraperitoneal injection. b. BLI images measuring tumor loads in experimental mice over time. c. Quantification of b (n = 5). d. Kaplan–Meier survival curves (n = 5). e. Studying the *in vivo* functionality of MAIT cells in the presence or absence of 5-OP-RU. FACS analyses of T cell activation marker CD69 and CD25 at Day 14 (n = 5). f-h. Studying the *in vivo* antitumor efficacy of MAIT cells in a subcutaneous liver cancer xenograft NSG mouse model. 5-OP-RU was administrated to experimental mice semi-weekly. f. Experimental design. s.c., subcutaneous injection. g. Measurement of tumor weight on day 34 (n = 5). h. Measurements of tumor size over time (n = 5). Representative of 3 experiments. Data are presented as the mean ± SEM. ns, not significant, **p < 0.01, ***p < 0.001, ****p < 0.0001 by Student’s t test (e), by one-way ANOVA (c, g), or by log rank (Mantel-Cox) test adjusted for multiple comparisons (d). See also Supplementary Fig. 5.

Mice were assigned to four treatment groups comprising adoptive transfer of MAIT cells alone, conventional T cells alone, MAIT cells with weekly intraperitoneal 5-OP-RU administration, and conventional T cells with 5-OP-RU (Fig. 5a). Bioluminescence imaging demonstrated progressive tumor expansion in vehicle, T cell, and T cell plus 5-OP-RU groups, indicating that conventional T cells lacked intrinsic tumor control capacity in this setting (Fig. 5b&5c). Adoptive MAIT cell transfer alone resulted in modest tumor suppression accompanied by a limited survival benefit (Fig. 5b–d). Strikingly, systemic administration of 5-OP-RU in MAIT-treated mice induced rapid and sustained tumor clearance, as evidenced by near-complete loss of hepatic BLI signal and prolonged survival (Fig. 5b–d). These results indicate that microbial metabolite delivery functions as a critical activating signal enabling effective MAIT antitumor activity *in vivo*.

Flow cytometric analysis of liver-resident human lymphocytes confirmed MAIT cell activation following adoptive transfer. 5-OP-RU administration was associated with increased expression of activation markers CD25 and CD69 on MAIT cells, consistent with in situ metabolite-driven activation (Fig. 5e).

To assess whether these findings extended to localized solid tumors, a complementary subcutaneous xenograft model was generated by implantation of SK-HEP-1-MR1-FG cells (Fig. 5f). Therapeutic groups mirrored those used in the orthotopic study, with semi-weekly subcutaneous 5-OP-RU delivery. Tumor growth kinetics revealed progressive tumor expansion in vehicle and T cell–treated cohorts, whereas MAIT cell transfer alone conferred partial growth delay (Fig. 5g&5h and Supplementary Fig. 5). In contrast, combined MAIT cell and 5-OP-RU treatment produced marked suppression of tumor growth, resulting in reduced tumor volume and tumor weight relative to all comparator groups (Fig. 5g&5h and Supplementary Fig. 5).

Collectively, these orthotopic and subcutaneous studies demonstrate that systemic microbial metabolite administration acts as a pharmacologic precision immune activator capable of licensing MAIT cells for potent tumor control across distinct *in vivo* contexts. These findings establish metabolite-driven MR1 signaling as a tractable therapeutic axis for harnessing endogenous MAIT biology in cancer immunotherapy.

### Microbial metabolite signaling drives cytotoxic state conversion in MAIT cells

To define the molecular basis of microbial metabolite–mediated MAIT activation, we performed single-cell RNA sequencing of MAIT cells cultured alone, co-cultured with tumor cells, or co-cultured with tumor cells in the presence of the microbial metabolite 5-OP-RU for 24 hours (Fig. 6a). This experimental design enabled resolution of MAIT cell state transitions induced by tumor encounter and microbial metabolite signaling within a shared transcriptional landscape.

**Fig. 6.**
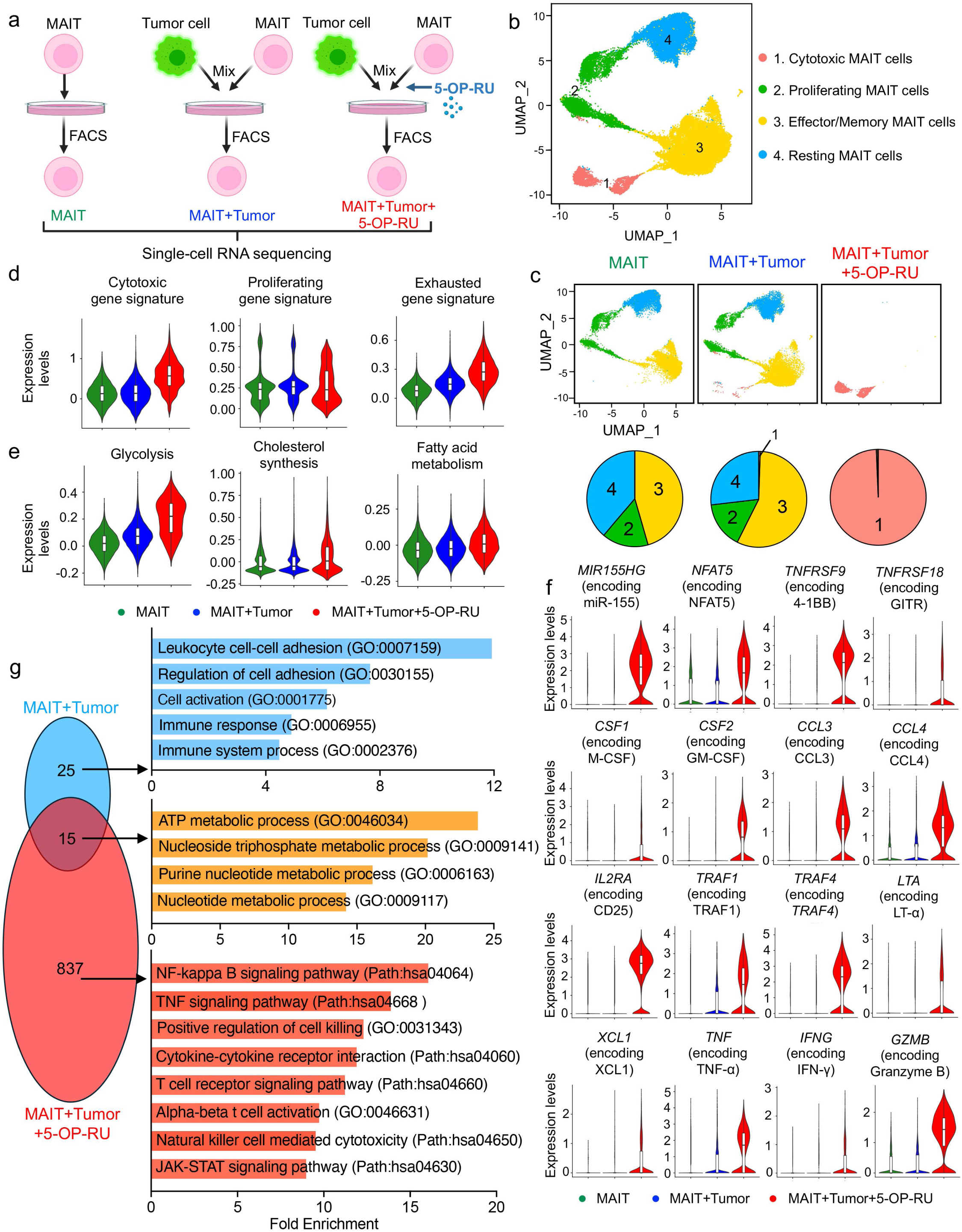
Microbial metabolite signaling drives cytotoxic state conversion in MAIT cells. a. Schematic showing the experimental design to study the gene profiling of MAIT cells using scRNA-seq. MAIT cells were cultured alone (MAIT), cocultured with tumor cells (MAIT+Tumor) or cocultured with tumor cells in the presence of 5-OP-RU (MAIT+Tumor+5-OP-RU). Cells were subsequently isolated by FACS and subjected to scRNA-seq analysis. b. Combined UMAP plot showing the formation of four major MAIT cell clusters: (1) Cytotoxic MAIT cells, (2) Proliferating MAIT cells, (3) Effector/memory MAIT cells, and (4) Resting MAIT cells. Each dot represents a single cell and is colored according to its cell cluster assignment. c. Individual UMAP plots showing cell cluster composition of the indicated samples, and and pie charts showing cell cluster proportions of the indicated samples. Each dot represents a single cell and is colored according to its cell cluster assignment. d. Violin plots showing the expression distribution of cytotoxic, proliferating and exhausted gene signatures in the indicated samples. e. Violin plots showing the expression levels of the metabolic pathways (glycolysis, cholesterol synthesis, and fatty acid metabolism) in the indicated samples. f. Violin plots showing the expression levels of representative pro-inflammatory genes in the indicated samples. g. Venn diagram illustrating the numbers of shared and unique differentially expressed genes (DEGs) of MAIT+Tumor and MAIT+Tumor+5-OP-RU samples relative to non–tumor-challenged MAIT cells. The pathway analyses were conducted using the shared and unique DEGs from each comparison. The indicated pathway in each category are shown in the bar plots. The experiment was performed once; cells collected from three repeated experiments were combined for analyses. In the violin plots (d, e, and f), box and whisker plots exhibit the minimum, lower quartile, median, upper quartile and maximum expression levels of each sample. See also Supplementary Fig. 6.

Uniform manifold approximation and projection (UMAP) of the integrated dataset revealed four transcriptionally distinct MAIT cell clusters (Fig. 6b). Based on established gene signatures, cluster 1 corresponded to cytotoxic MAIT cells, cluster 2 to proliferating MAIT cells, cluster 3 to effector/memory MAIT cells, and cluster 4 to resting MAIT cells (Supplementary Fig. 6)^62–65^. Notably, MAIT cells cultured alone or with tumor cells exhibited largely overlapping state distributions, predominantly occupying effector/memory, resting, and proliferative compartments (Fig. 6c). Tumor exposure alone induced only minimal cytotoxic conversion, with approximately 1% of cells adopting a cytotoxic phenotype. In contrast, addition of the microbial metabolite 5-OP-RU resulted in a striking state shift, with >99% of MAIT cells transitioning into the cytotoxic cluster (Fig. 6c), indicating metabolite-driven dominance of a cytotoxic transcriptional program.

Comparative gene signature analysis further supported this transition. Cytotoxic and exhaustion-associated gene programs were significantly upregulated following 5-OP-RU stimulation, whereas proliferative signatures remained largely unchanged (Fig. 6d), suggesting preferential induction of effector differentiation rather than proliferative expansion. Concomitantly, metabolic pathway analysis revealed increased expression of genes involved in glycolysis, cholesterol biosynthesis, and fatty acid metabolism (Fig. 6e), consistent with metabolic remodeling required to sustain heightened effector function.

Gene-level interrogation demonstrated robust induction of canonical cytotoxic and inflammatory mediators in the 5-OP-RU condition, including *MIR155HG*, *NFAT5*, *TNFRSF9*, *TNFRSF18*, *CSF1*, *CSF2*, *CCL3*, *CCL4*, *IL2RA*, *TRAF1*, *TRAF4*, *LTA*, *XCL1*, *TNF*, *IFNG*, and *GZMB* (Fig. 6f). These transcriptional changes collectively reflect coordinated activation of co-stimulatory signaling, cytokine production, and cytolytic machinery characteristic of highly functional effector lymphocytes.

Differential expression analysis further highlighted the magnitude of microbial metabolite–induced remodeling. Comparison of MAIT cells co-cultured with tumor cells versus MAIT cells alone identified 25 differentially expressed genes, whereas addition of 5-OP-RU yielded 837 differentially expressed genes, with 15 genes shared between conditions (Fig. 6g). Gene set enrichment analysis revealed that tumor co-culture alone primarily enriched pathways related to leukocyte adhesion, immune activation, and cell–cell interaction, whereas microbial metabolite exposure preferentially activated metabolic programs alongside effector immune pathways, including NF-κB signaling, TNF signaling, cytokine–cytokine receptor interaction, T cell receptor signaling, natural killer cell–mediated cytotoxicity, and JAK–STAT signaling (Fig. 6g).

Collectively, these single-cell analyses demonstrate that microbial metabolite signaling functions as a dominant regulator of MAIT cell fate, driving global transcriptional and metabolic reprogramming toward a cytotoxic effector state. These findings provide mechanistic evidence that metabolite availability governs MAIT functional differentiation within tumor contexts.

### Microbial metabolite enables MAIT-mediated elimination of immunosuppressive myeloid cells

The immunosuppressive TME represents a major barrier to effective immunotherapy in solid malignancies^66–69^, characterized by the accumulation of suppressive myeloid populations that promote tumor progression, facilitate immune evasion, and limit therapeutic responses^70–72^. Given the MR1-dependent cytotoxic capacity of metabolite-activated MAIT cells, we hypothesized that microbial metabolite signaling could enable MAIT cells to directly target immunosuppressive TME components and thereby remodel the immune landscape. To interrogate this possibility, we employed four complementary experimental systems spanning patient-derived samples, *in vitro* macrophage systems, 3D organoid cultures, and humanized *in vivo* models.

We first evaluated MAIT cell activity within primary liver tumor samples derived from patients (Fig. 7a). These specimens contained heterogeneous immune populations typical of the liver TME, including tumor-associated macrophages (TAMs), myeloid-derived suppressor cells (MDSCs), granulocytes, T cells, and B cells (Supplementary Fig. 7a). MR1 expression analysis revealed elevated MR1 levels on TAM and MDSC populations relative to other immune subsets (Supplementary Fig. 7b). Upon microbial metabolite activation with 5-OP-RU, MAIT cells selectively eliminated MR1^+^ TAM and MDSC populations while sparing T and B cells (Fig. 7b), indicating preferential targeting of immunosuppressive myeloid compartments within primary tumors.

**Fig. 7.**
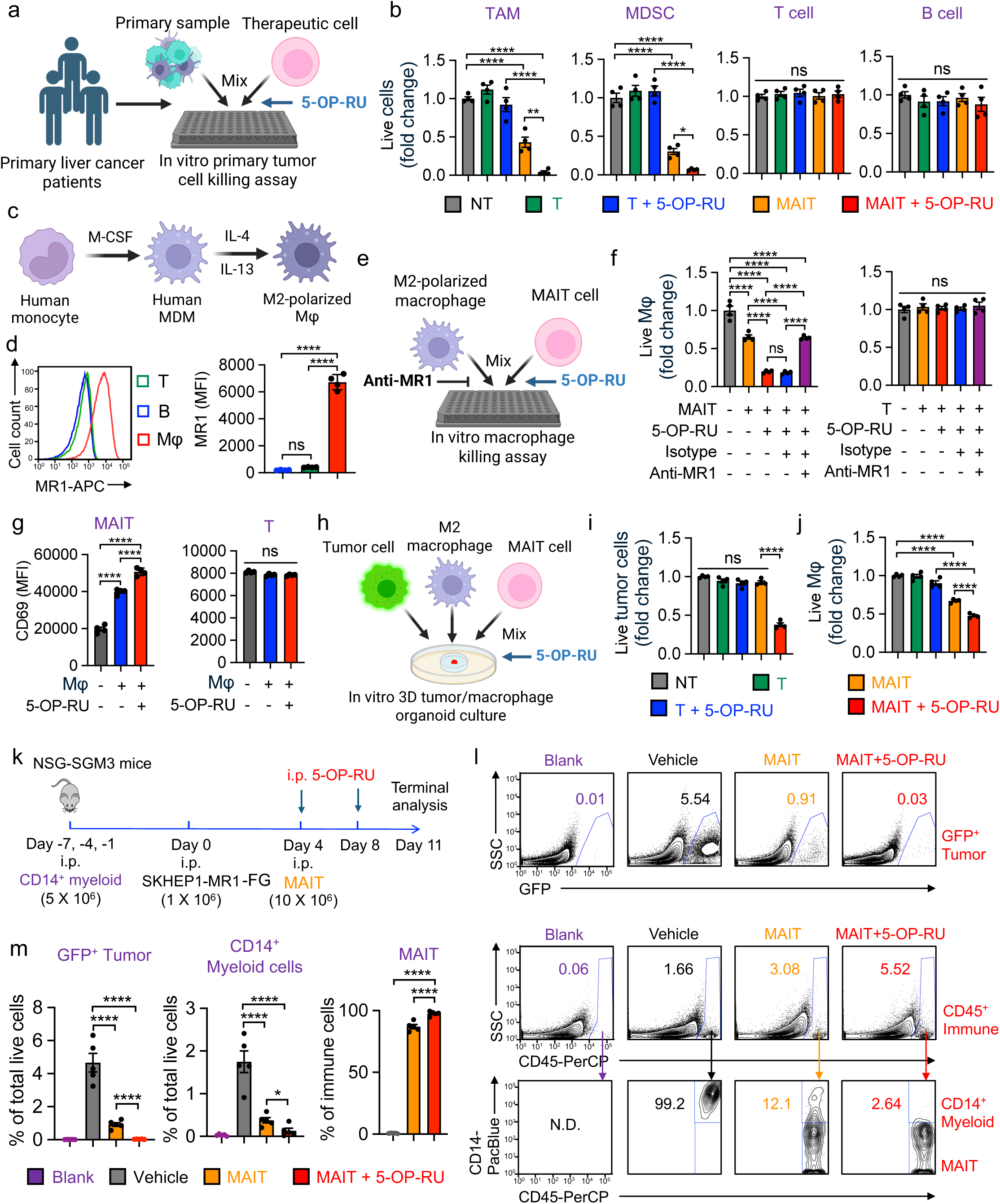
Microbial metabolite enables MAIT-mediated elimination of immunosuppressive myeloid cells. a-b. Studying microbial metabolite-mediated TME targeting by MAIT cells using primary liver cancer patient samples. Primary samples were cocultured with MAIT cells in the presence or absence of 5-OP-RU. T cells were included as a therapeutic cell control. Donor-matched patient T and B cells were included as target cell controls. a. Experimental design. b. TAM and MDSC killing data by MAIT cells at 24 h (n = 4). c-d. *In vitro* generation and polarization of human monocyte-derived M2 macrophages from healthy donor PBMCs. c. Experimental design. M-CSF, macrophage colony-stimulating factor; MDM, monocyte-derived macrophage; Mφ, macrophage. d. FACS analyses of MR1 on M2-polarized macrophages. Donor-matched T and B cells were included as controls. e-g. Studying microbial metabolite-mediated macrophage targeting by MAIT cells using an *in vitro* mixed macrophage/MAIT cell (Mφ/MAIT) reaction assay. M2-polarized macrophages were cocultured with MAIT cells in the presence or absence of 5-OP-RU. T cells were included as a therapeutic cell control. Anti-MR1 antibody was added into the coculture to block MAIT TCR recognition. e. Experimental design. f. M2-polarized macrophage killing data by MAIT cells at 24 h (n = 4). g. FACS detection of T cell activation marker CD69 on MAIT cells at 24 h (n = 4). h-j. Studying microbial metabolite-mediated TAM targeting by MAIT cells using an *ex vivo* 3D TME mimicry culture. M2-polarized macrophages and liver cancer cells were cocultured with MAIT cells in the presence or absence of 5-OP-RU. T cells were included as a therapeutic cell control. h. Experimental design. i. Tumor cell killing data by MAIT cells at 24 h (n = 4). j. M2-polarized macrophage killing data by MAIT cells at 24 h (n = 4). k-m. Studying microbial metabolite-mediated TME targeting by MAIT cells using humanized NSG-SGM3 myeloid cell–bearing xenograft model. k. Experimental design. l. FACS plots showing GFP⁺ liver tumor cells, human CD14+ myeloid cells, and MAIT cells in mouse peritoneal cavity collected from the indicated groups at day 11. m. Quantification of l (n = 5). Representative of 2 (k-n) and 3 (a-j) experiments. Data are presented as the mean ± SEM. ns, not significant, *p < 0.05, **p < 0.01, ***p < 0.001, ****p < 0.0001 by one-way ANOVA (b, d, f, g, i, j, m). Note the comparison between MAIT and MAIT + 5-OP-RU in m is analyzed by Student’s t test. See also Supplementary Fig. 7 and 8.

To mechanistically validate this observation, M2-polarized macrophages were generated from healthy donor monocytes as a model of suppressive macrophages (Fig. 7c). These cells exhibited high MR1 expression (Fig. 7d) and were efficiently eliminated by MAIT cells following 5-OP-RU stimulation (Fig. 7e&7f). Cytotoxicity was markedly reduced by MR1 blockade, confirming an MR1-dependent, MAIT TCR-mediated mechanism (Fig. 7f). Consistent with functional activation, CD69 expression increased selectively on MAIT cells but not conventional T cells following microbial metabolite exposure (Fig. 7g). Parallel experiments using 5-OE-RU similarly enhanced MAIT-mediated macrophage killing (Supplementary Fig. 8), indicating that this effect generalizes across MR1 ligands.

To assess MAIT function within an organized suppressive microenvironment, we generated 3D tumor organoids composed of liver tumor cells and M2 macrophages (Fig. 7h)^73,74^. In this context, conventional T cells exhibited neither antitumor nor anti-macrophage activity (Fig. 7i&7j). MAIT cells alone displayed limited tumor cytotoxicity; however, addition of 5-OP-RU restored robust MAIT tumor killing despite the presence of suppressive macrophages (Fig. 7i&7j). These findings suggest that microbial metabolite activation enables MAIT cells to overcome macrophage-mediated suppression, likely through selective depletion of immunosuppressive myeloid cells.

We next evaluated these interactions *in vivo* using NSG-SGM3 mice, which support human myeloid cell engraftment through expression of human SCF, GM-CSF, and IL-3^75,76^. Repeated administration of human CD14^+^ myeloid cells established a sustained suppressive compartment (Fig. 7k). Following tumor implantation and MAIT adoptive transfer, microbial metabolite administration significantly enhanced MAIT-mediated tumor control, as evidenced by reduced detection of GFP^+^ tumor cells within the peritoneal cavity (Fig. 7l&7m). Importantly, analysis of the TME revealed robust myeloid engraftment in control conditions, whereas MAIT cell transfer reduced these populations and microbial metabolite activation further amplified this effect, resulting in pronounced depletion of human myeloid cells (Fig. 7l&7m).

Collectively, these multi-scale analyses demonstrate that microbial metabolite signaling enables MAIT cells to selectively eliminate MR1-expressing immunosuppressive myeloid populations. These findings establish microbial metabolite as a pharmacologic precision immune activator that not only enhances MAIT tumor cytotoxicity but also actively remodels the TME, positioning microbial metabolite-driven MAIT activation as a strategy to overcome myeloid-mediated immune suppression in solid tumors.

## Discussion

In this study, we systematically define microbial metabolite–driven MR1 signaling as a tractable strategy to mobilize human MAIT cells for anti-tumor immunity across solid tumors. We first establish that circulating human MAIT cells possess a relatively uniform effector-memory phenotype and an innate-like cytotoxic transcriptional program compared with conventional T cells, indicating that these cells are intrinsically poised for rapid effector responses (Fig. 1). Building on this baseline state, we demonstrate that riboflavin-derived MR1 ligands function as pharmacologic precision immune activators capable of selectively mobilizing and expanding MAIT cells within physiologic immune environments derived from both healthy donors and cancer patients (Fig. 2). Functionally, microbial metabolite licensing converts MAIT cells from minimally active lymphocytes into potent tumor cytotoxic effectors across a broad panel of malignancies. Across ten human tumor cell lines spanning liver, ovarian, melanoma, lung, breast, and colorectal cancers, as well as primary patient-derived tumor cells, metabolite activation enables robust MR1-dependent tumor killing, establishing MR1 as a shared molecular interface through which MAIT cells can recognize tumors with heterogeneous baseline MR1 expression (Fig. 3). This activation is accompanied by coordinated effector and inflammatory responses while avoiding indiscriminate induction of tumor-supportive programs, indicating that microbial metabolite signaling drives a selective immune activation state rather than generalized immune perturbation (Fig. 4).

We further demonstrate the translational relevance of this pathway by showing that pharmacologic MR1 activation enables MAIT-mediated tumor control *in vivo* across both disseminated and localized tumor contexts using orthotopic and subcutaneous xenograft models, as well as humanized mouse systems that recapitulate human myeloid–tumor interactions (Fig. 5). Mechanistically, single-cell transcriptional profiling reveals that microbial metabolite exposure acts as a dominant regulatory signal driving a population-wide transition toward a cytotoxic MAIT cell state, providing a unifying explanation for the rapid functional activation observed across experimental systems (Fig. 6). Beyond direct tumor killing, metabolite-activated MAIT cells also remodel the TME by selectively eliminating MR1-expressing immunosuppressive myeloid populations, including tumor-associated macrophages and myeloid-derived suppressor cells (Fig. 7). Together, these findings establish microbial metabolite–MR1 signaling as a precision immune activation axis capable of simultaneously enabling direct tumor cytotoxicity and dismantling myeloid-mediated immune suppression.

From a broader conceptual perspective, our findings suggest that the principal limitation of MAIT cells in cancer is not intrinsic effector capacity but insufficient antigenic licensing within tumor environments. Human MAIT cells display a pre-armed effector-memory phenotype and innate-like cytotoxic programming, yet within tumors they often remain functionally restrained. By demonstrating that microbial metabolites are sufficient to trigger rapid cytotoxic state conversion, our study reframes MAIT cells as a latent cytotoxic lymphocyte population whose tumor reactivity can be governed by an MR1-ligand “on-switch.” This conceptual framework differs from conventional T cell immunotherapy paradigms that rely on tumor antigen specificity and instead positions MAIT cells as modular effector cells whose activity can be pharmacologically engaged through metabolite signaling.

Most mechanistic studies investigating microbial metabolite–mediated MAIT activation have been conducted in murine systems, and these studies have generally concluded that MR1 ligands alone are insufficient to drive robust MAIT responses *in vivo*. Instead, effective activation in mice typically requires additional inflammatory signals, such as co-administration of Toll-like receptor (TLR) agonists, and reported antitumor effects often depend in part on secondary activation of other immune populations, including NK cells^31,32,77^. These observations have led to the prevailing view that microbial metabolites function primarily as cooperative rather than autonomous activators of MAIT-mediated immunity. In contrast, our findings demonstrate that microbial metabolites alone are sufficient to directly license potent cytotoxic activity in human MAIT cells. This divergence likely reflects fundamental biological differences between murine and human MAIT compartments. Mouse MAIT cells are relatively rare, generally constituting less than 1% of T cells, and are predominantly composed of IL-17–producing MAIT17 subsets with limited representation of IFN-γ–producing MAIT1 populations^3,33,34,78^. By contrast, human MAIT cells are more abundant and exhibit broader functional plasticity characterized by co-expression of the transcriptional regulators RORγt and T-bet, enabling concurrent type-1 and type-17 effector programs^3,4,33,34^. These quantitative and qualitative differences suggest that murine models may emphasize indirect or cooperative immune mechanisms for MAIT-mediated tumor control, whereas human MAIT cells possess a more intrinsically cytotoxic effector program that can be directly engaged through MR1–metabolite signaling. Our results therefore highlight the importance of studying MAIT biology in human systems and suggest that microbial metabolite–based activation strategies may have greater therapeutic potential in humans than predicted from mouse studies.

In the future, strategies to augment MAIT cell abundance may be required in settings where endogenous MAIT frequencies are insufficient to achieve therapeutic benefit. Adoptive transfer approaches using *ex vivo*–expanded MAIT cells derived from autologous, allogeneic, or stem cell–based sources represent promising avenues to overcome this limitation^48,79,80^. Moreover, rather than administering defined microbial metabolites, exploitation of bacteria capable of producing MR1 ligands provides an alternative strategy to sustain MAIT activation in situ^52,81^. Engineering commensal or tumor-targeting microbial platforms to generate elevated levels of riboflavin-derived intermediates could persistently modulate MAIT cell activity within tissue microenvironments Programmable^82,83^. In parallel, advances in drug delivery technologies may enhance the therapeutic feasibility of microbial metabolite–based interventions^84^. Formulation strategies incorporating polymers, lipid nanoparticles, or other delivery systems could enable controlled release, improved tissue localization, and reduced systemic exposure of MR1 ligands^85,86^. Chemical engineering approaches aimed at stabilizing MAIT antigens, including optimized solvent formulations and structural modification, may further improve pharmacologic performance^87^. Together, these approaches outline a translational framework for harnessing microbial metabolite–MR1 signaling to unlock the therapeutic potential of MAIT cells in solid tumor immunotherapy.

## Materials and Methods

### Study approval

This study complies with all relevant ethical regulations. All experiments involving primary liver cancer patient samples were approved by the Ronald Reagan UCLA Medical Center (IRB# IRB-25-0948). Animal studies were approved by the Division of Laboratory Animal Medicine at UCLA. Healthy donor PBMCs were provided by the UCLA/CFAR Virology Core Laboratory without identification information under federal and state regulations.

### Mice

NOD.Cg-*Prkdcscid Il2rgtm1Wjl*/SzJ (NOD-*scid* IL2Rgnull, NSG, cat. no. #005557), and NOD.Cg-*Prkdcscid Il2rgtm1Wjl* Tg(CMV-IL3,CSF2,KITLG)1Eav/MloySzJ (NOD-scid IL2Rgnull-3/GM/SF,NSG-SGM3, cat. no. #013062) mice were purchased from The Jackson Laboratory, and maintained in animal facilities of the UCLA in a temperature-controlled environment (68 °F to 79 °F) with a 12-hour light cycle. 6- to10-week-old mice were used for all experiments. All mice were bred and maintained under specific pathogen-free conditions. All animal experiments were approved by the Institutional Animal Care and Use Committee (IACUC) of UCLA, and all animal procedures were conducted in accordance with the animal care and use regulations of the Division of Laboratory Animal Medicine (DLAM) at UCLA. Experimental mice were randomly assigned to treatment groups to avoid statistically significant differences in the baseline tumor burden. Both male and female mice were included in the study, and no significant sex-dependent differences in tumor growth or treatment response were observed in the tumor models used.

### Media and reagents

Recombinant human IL-2, IL-7, and IL-15 were purchased from PeproTech. Fetal bovine serum (FBS), and β-mercaptoethanol (β-ME) were purchased from Sigma. Penicillin-streptomycin-glutamine (P/S/G), MEM nonessential amino acids (NEAA), HEPES buffer solution, and sodium pyruvate were purchased from Gibco. Normocin was purchased from InvivoGen. The RPMI 1640 cell culture medium and the DMEM cell culture medium were purchased from Thermo Fisher Scientific. The CryoStor Cell Cryopreservation Media CS10 was purchased from MilliporeSigma. The C10 medium was made of RPMI 1640 cell culture medium supplemented with FBS (10% v/v), P/S/G (1% v/v), NEAA (1% v/v), HEPES (10 mM), sodium pyruvate (1 mM), β-ME (50 μM), and Normocin (100 μg/ml). The D10 medium was made of DMEM supplemented with FBS (10% v/v), P/S/G (1% v/v), and Normocin (100 μg/ml). The R10 medium was made of RPMI 1640 supplemented with FBS (10% v/v), P/S/G (1% v/v), and Normocin (100 μg/ml). 5-Amino-4-D-ribitylaminouracil Dihydrochloride (5-A-RU; 90%; cat. no. TRC-A629245) was purchased from TorontoResearchChemicals. Methylglyoxal (MGO; cat. no. M0252-25ML) and glyoxal (GO; cat. no. 50649-100ML) solution was purchased from Sigma. 5-OP-RU was generated by mixing 5-A-RU (1 μM) with excess MGO (0.1%) to allow rapid and saturating formation of the 5-OP-RU ligand. 5-OE-RU was generated by mixing 5-A-RU (1 μM) with excess GO (0.1%) to allow rapid and saturating formation of the 5-OE-RU ligand. Both ligand mixtures were prepared immediately prior to use and applied freshly in subsequent experiments.

### Lentiviral vectors

All lentiviral vectors used in this study were constructed from a parental vector pMNDW^88,89^. The 2A sequence derived from foot-and-mouth disease virus (F2A) was used to link the inserted genes to achieve co-expression. The Lenti/FG vector was constructed by inserting a synthetic bicistronic gene encoding Fluc-P2A-EGFP into the pMNDW ^88,89^. The Lenti/MR1 vector was constructed by inserting a synthetic gene encoding human MR1 into the pMNDW. The synthetic gene fragments were obtained from GenScript and IDT. The Lenti/CRISPRv2-MR1 vector was purchased from GenScript (GenCRISPR gRNA Construct), containing the U6 promoter–driven gRNA scaffold, Cas9 nuclease, and puromycin selection cassette^90^. The gRNA sequence used to target human MR1 is GGATGGGATCCGAAACGCCC (5′–3′)^91^. The Lenti/CRISPRv2-MR1 vector was purchased from Genscript with gRNA GGATGGGATCCGAAACGCCC (5’-3’). Lentiviruses were produced using human embryonic kidney 293T (HEK293T) cells (ATCC), following a standard transfection protocol using the Trans-IT-Lenti Transfection Reagent (Mirus Bio) and a centrifugation concentration protocol using the Amicon Ultra Centrifugal Filter Units, according to the manufacturer’s instructions (MilliporeSigma).

### Stable tumor cell lines

Human liver cancer cell lines HepG2 (cat. no. HB-8065), Hep3B (cat. no. HB-8064), SNU-423 (cat. no. CRL-2238), and SK-HEP-1 (cat. no. HTB-52), melanoma cell line A375 (cat. no. CRL-1619), breast cancer cell line MDA-MB-231 (cat. no. HTB-26), colorectal cell line HCT116 (cat. no. CCL-247), and ovarian cancer cell lines OVCAR3 (cat. no. HTB-161), and SKOV3 (cat. no. HTB-77) were purchased from the ATCC. HEP3B and SNU423 are human HCC cell lines, HEPG2 is a human hepatoblastoma cell line, and SKHEP1 is a human endothelial-like liver cancer cell line. In this study, SK-HEP-1 cells were employed to generate an orthotopic liver tumor model due to their rapid migration and robust growth within the murine liver.

The parental tumor cell lines were transduced with Lenti/FG vector to produce stable tumor cell lines overexpressing FG. SK-HEP-1-FG cell line was further transduced with Lenti/MR1 to produce stable tumor cell lines overexpressing human MR1. SK-HEP-1-FG was also transduced with Lenti/CRISPRv2-MR1 to knock out human MR1 expression. 72 hours post lentivector transduction, cells were subjected to flow cytometry sorting and/or puromycin selection to isolate gene-engineered cells for generating stable cell lines. Twelve stable tumor cell lines were generated for this study, including HepG2-FG, Hep3B-FG, SNU423-FG, SK-HEP-1-FG, SK-HEP-1-MR1-FG, SK-HEP-1-FG^MR1-/-^, A375-FG, MDA-MB-231-FG, HCT116-FG, OVCAR3-FG, SKOV3-FG cell lines. All tumor cell lines utilized in this study underwent short tandem repeat (STR) profiling, and the resulting profiles were compared to established databases to confirm accurate identification. Furthermore, the cell lines were regularly screened for mycoplasma contamination to preserve their integrity and authenticity.

### Liver and PBMC sample collection from liver cancer patients

Primary liver cancer patient samples, including liver tumor and peripheral blood samples, were collected at the Ronald Reagan UCLA Medical Center from consented patients through an IRB-approved protocol (IRB-25-0948) and processed. Information regarding the patients’ gender and age was not provided in this study to avoid including three or more indirect identifiers for the study participants (Supplementary Table 1). Patient gender was not considered in the study design and was determined based on self-reporting.

### PBMC collection from healthy donors

Healthy donor PBMCs were provided by the UCLA/CFAR Virology Core Laboratory without identification information under federal and state regulations. PBMCs were cryopreserved in Cryostor CS10 (Sigma St. Louis, MO, USA) using CoolCell (BioCision, Larkspur, CA, UCA), and were frozen in liquid nitrogen for storage and to supply all experiments.

### Antibodies and flow cytometry

Fluorochrome-conjugated antibodies specific for human CD3 (clone HIT3a, Pacific Blue, PerCP, or PE-conjugated, 1:500; cat. no. 300330), CD4 (clone OKT4, PE-Cy7, PerCP, or FITC-conjugated, 1:500; cat. no. 317432), CD8 (clone SK1, PE, APC-Cy7, or APC-conjugated, 1:300, cat. no. 344706), CD45 (clone H130, PerCP, PE, or Pacific Blue-conjugated, 1:500; cat. no. 304026), CD56 (clone QA18A21, PE-Cy7, FITC, or PerCP-conjugated, 1:10; cat. no. 318342), CD69 (clone FN50, PE-Cy7, FITC or PerCP-conjugated, 1:50; cat. no. 310912), MR1 (clone 26.5, PE or APC-conjugated,1:50; cat. no. 361108), TCR Vα7.2 (clone 3C10, APC-Cy7, PE, or APC-conjugated,1:50; cat. no. 351708), IFN-γ (clone B27, PE-Cy7-conjugated, 1:50; cat. no. 506518), Granzyme B (clone QA16A02, APC-conjugated, 1:2,000; cat. no. 372204), Perforin (clone dG9, PE-Cy7-conjugated, 1:50 or 1:100; cat. no. 308126), IL-2 (clone MQ1-17H12, APC-Cy7-conjugated, 1:50; cat. no. 500342), IL-17A (clone BL168, APC-conjugated, 1:50; cat. no. 512333), T-bet (clone QA18A24, PE-Cy7-conjugated, 1:50; cat. no. 644824), CD161 (clone HP-3G10, PerCP, FITC or APC-conjugated, 1:50; cat. no. 339934), CD25 (clone BC96, APC-Cy7, or PE-conjugated, 1:200; cat. no. 356104), CD14 (clone HCD14, PacBlue-conjugated, 1:100; cat. no. 367122), CD11b (clone ICRF44, FITC-conjugated, 1:500; cat. no. 101206), CD19 (clone SJ25C1, APC or APC-Cy7-conjugated, 1:200; cat. no. 302218), CD20 (clone 2H7, PE/Cyanine7-conjugated, 1:200; cat. no. 302312), CD206 (clone 15-2, APC-Cy7 or APC-conjugated, 1:200; cat. no. 321110), CD107a (clone H4A3, PE-Cy7-conjugated, 1:200; cat. no. 328618), CD62L (clone DREG-56, APC-Cy7 or APC-conjugated, 1:200; cat. no. 304822) and CD45RO (clone UCHL1, PE-Cy7 or APC-Cy7-conjugated, 1:100; cat. no. 304230) were purchased from BioLegend. Fluorochrome-conjugated antibody specific for human PLZF (clone # 6318100, Alexa Fluor® 488-conjugated, 1:50; cat. no. IC2944G-100UG) was purchased from R&D Systems. Fluorochrome-conjugated antibody specific for human RORγt (clone Q21-559, PE-conjugated, 1:50; cat. no. 563081) was purchased from BD Biosciences. Fixable Viability Dye eFluor506 (e506, 1:500; cat. no. 65-0866-18) was purchased from Affymetrix eBioscience. Mouse Fc Block (anti-mouse CD16/32; cat. no. 553142) was purchased from BD Biosciences, and human Fc Receptor Blocking Solution (TrueStain FcX; cat. no. 422302) was purchased from BioLegend. PE or APC-conjugated human MR1/5-OP-RU Tetramer (1:500 dilution) was obtained from NIH Tetramer Core Facility. All flow cytometry staining was performed following standard protocols, as well as specific instructions provided by the manufacturer of a particular antibody. Stained cells were analyzed using a MACSQuant Analyzer 10 flow cytometer (Miltenyi Biotech), following the manufacturers’ instructions. FlowJo software version 9 (BD Biosciences) was used for data analysis. For intracellular cytokine staining, cells were thawed and resuspended in C10 medium. Cells were stimulated with eBioscience™ Cell Stimulation Cocktail (Invitrogen/Thermo Fisher Scientific, cat. no. 00-4970-03; 1:500 dilution) in the presence of GolgiStop (BD Biosciences, cat. no. 554724; 1.5 μL/mL) to inhibit cytokine secretion and incubated for 6 h at 37 °C. Following stimulation, intracellular cytokine staining was performed using the BD Cytofix/Cytoperm Fixation/Permeabilization Kit (BD Biosciences, cat. no. 554714) according to the manufacturer’s instructions. For transcription factor staining, cells were stained using the eBioscience™ Foxp3 / Transcription Factor Staining Buffer Set (Thermo Fisher Scientific, cat. no. 00-5523-00) according to the manufacturer’s protocol. In our study, note the use of antibodies with identical clones but differing conjugated fluorochromes, with one typical antibody listed here.

### Luminex assay

MAIT cells were co-cultured with SK-HEP-1 tumor cells for 24 hours, after which culture supernatants were collected. Cytokine concentrations in the supernatants were quantified using a 38-plex human cytokine Luminex multibead array (Millipore Sigma), including GM-CSF, IL-1α, IL-2, IFN-γ, TNF-α, IL-17A, IL-17F, CCL4, CCL5 (RANTES), LT-α, IL-6, IL-8 (CXCL8), PDGF-AA, PDGF-AB/BB, and VEGF-A. Samples were acquired on a Luminex 200 System (Luminex), and analyte concentrations (pg/ml) were determined using standard curves generated for each cytokine. All Luminex assays and data analyses were performed by the UCLA Immune Assessment Core.

### Generation of MAIT cells from healthy donor or liver cancer patient PBMCs

PBMCs from healthy donors or liver cancer patients were used to generate MAIT cells. Cells were enriched via a two-step MACS protocol, first stained with PE-conjugated MR1/5-OP-RU tetramer and then labeled with Anti-PE MicroBeads (Miltenyi Biotec) for magnetic separation. The sorted MAIT cells were co-cultured with irradiated autologous PBMCs at a 1:5 ratio in C10 medium supplemented with 5-OP-RU (100 nM) and human IL-7 and IL-15 (10 ng/mL each). Cultures were maintained for 2 weeks with periodic IL-7 and IL-15 cytokine supplementation. Expanded MAIT cells could be subsequently purified by FACS to isolate MR1/5-OP-RU tetramer^+^TCR Vα7.2^+^CD3^+^ cells for downstream applications.

### Generation of *Escherichia coli* (*E. Coli*) supernatant

*Escherichia coli* K-12 MG1655 (purchased from ATCC; cat. no. 50-238-3578) was cultured overnight in LB broth at 37 °C with shaking. Bacterial growth was monitored by measuring optical density at 600 nm (OD600), and a standard curve was generated to estimate cell density. Cultures were then centrifuged to pellet bacteria, and the supernatants were collected and passed through a 0.2 µm membrane filter to obtain sterile filtrates. Filtered supernatants were aliquoted and stored at −80 °C for downstream assays.

### Generation of Human M2-Polarized Macrophage

Healthy donor PBMC-derived, M2-polarized macrophages were used in this assay. PBMCs were resuspended in serum-free RPMI 1640 medium (Corning Cellgro, cat. no. 10-040-CV) at 1 x 10^7^ cells/ml, plated in 10 cm dishes (10–15 ml per dish), and incubated at 37 °C with 5% CO_2_ for 1 h. Non-adherent cells were removed, and adherent monocytes were utilized to generate M2-polarized macrophages. To generate M2-polarized macrophages, monocytes were cultured in C10 medium supplemented with recombinant human M-CSF (10 ng/ml, Peprotech, cat. no. 300 − 25) for 6 days. On day 6, macrophages were detached using 0.25% Trypsin/EDTA (Gibco, cat. no. 25200-056), collected, and reseeded in 6- or 12-well plates (0.5-1 x 10^6^ cells/ml) for another 48 h with recombinant human IL-4 (10 ng/mL, Peprotech, cat. no. 214 − 14) and IL-13 (10 ng/ml, Peprotech, cat. no. 214 − 13) to induce M2 polarization. M2-polarized macrophages were then collected and used for flow cytometry.

### *In vitro* MAIT cell stimulation assay

MAIT cells were stimulated from either healthy donor or cancer patient PBMCs under the indicated conditions. A total of 10 x 10^6^ live cells were cultured in 5 mL C10 medium supplemented with human IL-7 and IL-15 (10 ng/mL each). Depending on the experimental setup, cultures were additionally treated with 5-OP-RU (100 nM) or 5-OE-RU (200 nM). The frequency and phenotype of MAIT cells (identified as CD3^+^MR1/5-OP-RU tetramer^+^TCR Vα7.2^+^ cells) were analyzed by flow cytometry.

### *In vitro* tumor cell killing assay

Tumor cells (1 x 10^4^ cells per well) were co-cultured with MAIT or T cells (at ratios indicated in the figures or figure legends) in Corning 96-well clear bottom black plates for 24 h in C10 medium. D-luciferin (150 mg/ml, Caliper Life Science) was added to cell cultures to quantify live tumor cells and luciferase activities were read out using an Infinite M1000 microplate reader (Tecan). Depending on the experimental condition, cultures were supplemented with 5-OP-RU (100 nM), 5-OE-RU (200 nM) or *E. Coli* culture supernatant as specified in the figure legends. In some experiments, MAIT or T cells after 24 h of tumor cell coculture were collected for phenotype and functional analyses.

Additional tumor killing experiments were conducted using SK-HEP-1-MR1-FG tumor cells co-cultured with MAIT cells. To study the MR1-mediated tumor cell killing mechanism, 10 μg/mL LEAF^TM^ purified anti-human/mouse/rat MR1 antibody (Clone 26.5, BioLegend, San Diego, CA, USA) or LEAF^TM^ purified mouse IgG2aκ isotype control antibody (Clone MOPC-173, BioLegend, San Diego, CA, USA) were added to co-cultures.

### *In vitro* assays using healthy donor PBMC samples

Healthy donor PBMC samples were analyzed using flow cytometry. T cells were identified as CD45^+^CD3^+^ cells, CD4 T cells were identified as CD4^+^ T cells, CD8 T cells were identified as CD8^+^ T cells, MAIT cells were identified as MR1/5-OP-RU tetramer^+^TCR Vα7.2^+^ T cells.

### *In vitro* assays using primary liver cancer patient samples

In one assay, the primary liver cancer patient samples were analyzed for tumor cell phenotype and the TME composition using flow cytometry. Liver tumor cells were sorted using a Human Tumor Cell Isolation Kit (Miltenyi Biotec) and/or identified as CD45^-^CD31^-^FAP (fibroblast activation protein)^-^ cells^92–94^, T cells were identified as CD45^+^CD3^+^ cells, B cells were identified as CD45^+^CD19^+^ or CD45^+^CD20^+^ cells, NK cells were identified as CD45^+^CD56^+^CD3^-^ cells, tumor associated macrophages (TAMs) were identified as CD45^+^CD11b^+^CD14^+^CD206^+^ cells, myeloid-derived suppressor cells (MDSCs) were identified as CD45^+^CD11b^+^CD14^-^CD206^med/low^ cells, granulocytes were identified as CD45^+^SSC^+^ cells. Surface expression of MR1 on tumor or/and immune cells were also analyzed using flow cytometry.

In another assay, the primary liver cancer patient samples were used to study tumor and TME cell killing by MAIT cells under various conditions. Patient samples (containing 1 x 10^5^ cells) were directly co-cultured with MAIT or T cells (5 x 10^5^ cells) in C10 medium in Corning 96-well Round Bottom Cell Culture plates for 24 h. Depending on the experimental condition, cultures were supplemented with 5-OP-RU (100 nM) as specified in the figure legends. At the end of culture, cells were collected, and the tumor and TME cell targeting by MAIT cells was assessed using flow cytometry by quantifying live human tumor cells (identified as MR1/5-OP-RU tetramer^-^CD45^-^ cells), TAMs (identified as CD45^+^CD11b^+^CD14^+^CD206^+^ cells), MDSCs (identified as CD45^+^CD11b^+^CD14^-^CD206^med/low^), T cells (identified as MR1/5-OP-RU tetramer^-^CD45^+^CD3^+^ cells), B cells (identified as MR1/5-OP-RU tetramer^-^CD45^+^CD3^-^CD19^+^ cells or MR1/5-OP-RU tetramer^-^CD45^+^CD3^-^CD20^+^ cells), and NK cells (identified as MR1/5-OP-RU tetramer^-^CD45^+^CD3^-^CD56^+^ cells). A total of 3 primary liver cancer patient samples were included in this assay.

### *In vitro* mixed Mφ/T reaction assay

PBMC-derived M2-polarized macrophages were cocultured with donor-mismatched T or MAIT cells at a ratio of 1:2 in 96-well round-bottom plates in C10 medium for 24 h at the presence or absence of 5-OP-RU (100 nM). At the end of cell culture, these cells were collected to study surface marker expression using flow cytometry. 10 μg/mL LEAF^TM^ purified anti-human/mouse/rat MR1 antibody (Clone 26.5, BioLegend, San Diego, CA, USA) or LEAF^TM^ purified mouse IgG2aκ isotype control antibody (Clone MOPC-173, BioLegend, San Diego, CA, USA) was added to co-cultures to study MR1-mediated macrophage killing mechanism.

### *In vitro* 3D tumor organoid targeting assay

To generate tumor organoids, a mixture of 2 x 10^5^ SK-HEP-1-MR1-FG tumor cells, 2 x 10^5^ M2 macrophages, and 2 x 10^6^ therapeutic cells (i.e., T or MAIT cells) were mixed in C10 medium. Mixed cells were centrifuged, resuspended in C10 medium at a concentration of 1 x 10^5^ cells/µl, adjusted to 5–10 μL per aggregate, and gently transferred onto a microporous membrane cell inserts (EMD Millipore, MA, USA, cat. no. PICM0RG50) to form a 3D human tumor/TAM/T-cell organoid in 6-well plates containing 1 ml of C10 medium per well^74,95^. Co-culture was maintained at the presence or absence of 5-OP-RU (100 nM) for 48 h. Following the co-culture period, organoids were mechanically dissociated with a 1 ml pipette and passed through a 70-µm nylon strainer to generate single-cell suspensions for downstream flow cytometry analysis.

### Single cell RNA sequencing (scRNA-seq)

In one analysis, publicly available scRNA-seq data were used to examine gene expression profiles of healthy donor PBMCs. The dataset was obtained from Synapse (accession code syn49637038)^39^. γδ T cells were excluded prior to analysis. Data processing, clustering, and annotation were performed using Seurat (v4.0.0) in R following the guidelines^96–98^. Briefly, after filtering the low-quality cells, the dataset was normalized using NormalizeData function, followed by selecting variable features across datasets using FindVariableFeatures functions. The dataset was then subjected to standard Seurat workflow for dimension reduction and clustering. Clusters of therapeutic cells were manually merged and annotated based on gene signatures reported from previous studies^64,65,99–104^. AddModuleScore was used to calculate module scores of each list of gene signatures, and FeaturePlot function was used to visualize the expression of each signature in the UMAP plots. VlnPlot function was used to assess gene expression differences between groups.

In a separate study, scRNA-seq was performed to characterize MAIT cell transcriptional responses following tumor cell coculture. The experimental design is illustrated in Fig. 6a. MAIT cells were cocultured with SK-HEP-1-MR1-FG liver tumor cells at the presence or absence of 5-OP-RU (100 nM) for 24 hours. The subsequent MAIT cells were FACS sorted and delivered to the UCLA TCGB Core for library construction and scRNA-seq.

Cells were quantified using a Cell Countess II automated cell counter (Invitrogen/ Thermo Fisher Scientific). A total of 10,000 cells from each experimental group were loaded on the Chromium platform (10X Genomics), and libraries were constructed using the Chromium Next GEM Single Cell 3′ Kit v3.1 and the Chromium Next GEM Chip G Single Cell Kit (10X Genomics), according to the manufacturer’s instructions. Library quality was assessed using the D1000 ScreenTape on a 4200 TapeStation System (Agilent Technologies). Libraries were sequenced on an Illumina NovaSeq using the NovaSeq S4 Reagent Kit (100 cycles; Illumina). Raw sequencing data were processed with Cell Ranger to generate gene expression matrices, which were subsequently analyzed in Seurat (v4.0.0) using the workflow described above.

### *In vivo* bioluminescence imaging (BLI)

BLI was performed using a Spectral Advanced Molecular Imaging HTX system (Spectral Instrument Imaging). Live animal images were captured 5 minutes after intraperitoneal (i.p.) injection of D-Luciferin (1 mg per 100 μL PBS per mouse) to obtain total body bioluminescence. The imaging data were analyzed using AURA imaging software (version 3.2.0, Spectral Instrument Imaging).

### *In vivo* antitumor efficacy study of MAIT cells: human orthotopic liver cancer xenograft NSG mouse model

Experimental design is shown in Fig. 5a. Briefly, on Day 0, NSG mice received i.v. inoculation of SK-HEP-1-MR1-FG human liver cancer cells (5 x 10^5^ cells per mouse). On Day 7, mice were randomized into five treatment groups: Vehicle control (100 μl PBS per mouse), T cells (1 x 10⁷ cells in 100 μl PBS per mouse), MAIT cells (1 x 10⁷ cells in 100 μl PBS per mouse), T cells plus 5-OP-RU (100 nM in 100 µL PBS), and MAIT cells plus 5-OP-RU (100 nM in 100 µL PBS). For microbial metabolite-treated groups, 5-OP-RU was administered i.p. weekly. Throughout the study, mice were monitored for survival, and tumor burden was assessed twice weekly using bioluminescence imaging (BLI). In a parallel study, mice were euthanized 7 days post MAIT or T cell infusion, and their tissues were collected for MAIT cell phenotype analysis by flow cytometry.

### *In vivo* antitumor efficacy study of MAIT cells: human subcutaneous liver cancer xenograft NSG mouse model

Experimental design is shown in Fig. 5f. Briefly, on Day 0, NSG mice received s.c. inoculation of SK-HEP-1-MR1-FG human liver cancer cells (5 x 10^5^ cells per mouse). On Day 7, mice were randomized into five treatment groups: Vehicle control (100 μl PBS per mouse), T cells (1 x 10⁷ cells in 100 μl PBS per mouse), MAIT cells (1 x 10⁷ cells in 100 μl PBS per mouse), T cells plus 5-OP-RU (100 nM in 100 µL PBS), and MAIT cells plus 5-OP-RU (100 nM in 100 µL PBS). For microbial metabolite-treated groups, 5-OP-RU was administered s.c. semi-weekly. Throughout the study, mice were monitored for survival, and tumor size was measured twice weekly using a Fisherbrand Traceable digital caliper (Thermo Fisher Scientific); tumor volumes were calculated by formula 1/2 × L × W^2^.

### *In vivo* antitumor and anti-TME efficacy study of MAIT cells: human liver cancer xenograft NSG-SGM3 mouse model

Experimental design is shown in Fig. 7k. Briefly, on Day -7, -4, and -1, NSG-SGM3 mice received i.p. injection of healthy donor PBMC-derived CD14^+^ myeloid cells (5 x 10^6^ cells per mouse) to establish a liver cancer TME enriched with TAM-like cells. On Day 0, NSG-SGM3 mice received i.p. inoculation of SK-HEP-1-MR1-FG human liver cancer cells (1 x 10^6^ cells per mouse). On Day 4, mice were randomized into three treatment groups: Vehicle control, MAIT cells, and MAIT cells plus 5-OP-RU. Mice in the Vehicle group received an intraperitoneal (i.p.) injection of PBS (100 μl per mouse). Mice in the MAIT and MAIT + 5-OP-RU groups received human MAIT cells (1 x 10⁷ cells in 100 μl PBS per mouse, i.p.). Subsequently, mice in the MAIT + 5-OP-RU group received i.p. injections of 5-OP-RU (100 nM in 100 μl PBS) on Days 5 and 8, whereas mice in the MAIT group received PBS vehicle injections at the same time points. At Day 11, mice were euthanized, and their peritoneal fluid were collected for tumor cell, human myeloid cell and MAIT cell phenotype analyses by flow cytometry. Blank NSG-SGM3 mice were included as a control.

### Statistics

Statistical data analysis was performed using GraphPad Prism 8 software (GraphPad). Two-tailed student’s t test was employed for pairwise comparisons. Ordinary one ANOVA followed by Tukey’s multiple comparisons test was used for multiple comparisons. Log-rank (Mantel-Cox) test adjusted for multiple comparisons was used for Kaplan–Meier survival curves analysis. Data are expressed as the mean ±SEM, unless otherwise indicated. In all figures and figure legends, n denotes the number of samples or animals utilized in the indicated experiments. A p-value of less than 0.05 was considered significant; ns indicates not significant; *p < 0.05, **p < 0.01, ***p < 0.001, ****p < 0.0001.

## Acknowledgements

We thank the University of California, Los Angeles (UCLA) animal facility for providing animal support; the UCLA Translational Pathology Core Laboratory (TPCL) for providing histology support; the UCLA Technology Centre for Genomics & Bioinformatics (TCGB) facility for providing RNA-seq services; the UCLA CFAR Virology Core for providing human cells; and the UCLA BSCRC Flow Cytometry Core Facility for cell sorting support; the UCLA Immune Assessment Core (IAC) for providing Luminex assays and analyses. We thank NIH Tetramer Core Facility for providing the tetramers. Some Figures were created with BioRender (https://biorender.com).

## AUTHOR CONTRIBUTIONS

Y.Z., Y-R.L., and L.Y. designed the experiments, analyzed the data, and wrote the manuscript. L.Y. and Y-R.L. conceived and oversaw the study, with assistance from Y.Z.. Y.Z. performed all experiments, with assistance from X.S., Y.C., N.M., C.Z., A.S.Z., Y.T., J.H., S.L., and A.W.. S.G. and V.G.A. provided the primary patient samples.

## Data Availability Statement

The scRNA-seq data generated in this study have been deposited in the public repository Gene Expression Omnibus Database: GSE322543. The following secure token has been created for review: czifiukynpmnjwh.

